# A Novel FNDC1-NAMPT-NAD axis is Implicated in Small and Large-vessel Arterial Disease and Drives Vascular Calcification

**DOI:** 10.1101/2025.07.08.663801

**Authors:** Sujin Lee, Yugene Guo, Lova P. Kajuluri, Kangsan Roh, Kuldeep Singh, Wanlin Jiang, Elizabeth Moore, Helena Tattersfield, Katrina Ostrom, Claire Birchenough, Adam Lee Johnson, Scott Krinsky, Houda Bouchouari, Mohammed Mohammedeh, Laurel Y. Lee, Gregory A. Wyant, Stephen S. Rich, Jerome L. Rotter, Kent D. Taylor, Shaista Malik, Robert Gerszten, Daniel Cruz, Russell P. Tracy, Hui Wu, Hua Zhang, Haojie Lu, Maryam Kavousi, Cathy Shanahan, Melinda Duer, Rafael Kramann, Rosalynn M. Nazarian, Matthew J. Eagleton, Yabing Chen, Clint L. Miller, Sagar U. Nigwekar, Rajeev Malhotra

## Abstract

Vascular calcification represents a convergent pathological feature of diverse cardiovascular diseases, yet the upstream molecular programs orchestrating this process remain poorly defined. Here, we uncover *fibronectin type III domain-containing* 1 (FNDC1) as a previously unrecognized regulator of vascular calcification across both microvascular and macrovascular beds. Integrative transcriptomic profiling of human calciphylaxis lesions and atherosclerotic coronaries identified FNDC1 as one of the most significantly upregulated genes. In primary human vascular smooth muscle cells, FNDC1 drove osteogenic phenotype switch and vascular calcification through activation of PI3K/AKT signaling and metabolic reprogramming. Mechanistically, FNDC1 directly binds to nicotinamide phosphoribosyltransferase (NAMPT) resulting in elevated intracellular NAD⁺ levels, thus coupling vascular signaling to control of NAD⁺ biosynthesis. In murine models, genetic deletion of *Fndc1* or pharmacologic inhibition of NAMPT suppressed arterial calcification and prolonged survival. Clinically, circulating FNDC1 levels were elevated in patients with both calciphylaxis and coronary artery disease and independently predicted cardiovascular risk in 42,687 UK Biobank participants. Together, these findings establish FNDC1 as a central mediator of vascular pathology and highlight the FNDC1– NAMPT-NAD^+^ axis as a promising target for therapeutic intervention.

## INTRODUCTION

Vascular calcification is a hallmark of atherosclerotic cardiovascular disease, the leading cause of morbidity and mortality worldwide.^1–3^ While macrovascular and microvascular calcifications have historically been considered distinct pathological processes, emerging evidence highlights a shared cellular and molecular framework-most notably, vascular smooth muscle cell (VSMC) osteogenic phenotype switch as a central driver of calcific remodeling.^4,5^

Calciphylaxis, a rare yet devastating syndrome characterized by rapid medial arteriolar calcification and downstream ischemic tissue necrosis, exemplifies the clinical severity of microvascular calcification, with mortality rates exceeding 50-80% within one year of diagnosis.^6–8^ Although associated with end-stage renal disease and metabolic derangements such as hyperphosphatemia and diabetes, the molecular drivers of calciphylaxis remain poorly understood, and no effective targeted therapies are available.^6,9^ Similarly, in coronary artery disease, large-vessel calcification contributes to plaque instability and is associated with adverse events, yet mechanistic insight has not translated into anti-calcific drugs.^10–12^

Medial calcification in calciphylaxis and intimal atherosclerotic calcification are associated with a phenotypic transition of VSMCs from a contractile to an osteogenic state, which is orchestrated by osteogenic transcription factors such as runt-related transcriptional factor-2 (RUNX2) and by metabolic rewiring that provides the energetic and biosynthetic substrates required for mineral deposition.^4,5,13^ These observations raise the possibility that macrovascular and microvascular calcification share conserved regulatory networks, despite occurring in anatomically and clinically distinct vascular beds.^10–12,14–16^

Here, we combined bulk and single-cell transcriptomic profiling of human calciphylaxis and coronary artery disease (CAD) tissues with gain and loss-of-function studies in VSMCs and mouse models to test the hypothesis that common signaling pathways govern calcification across vascular beds. We show that FNDC1 is a hub gene upregulated in both micro- and macrovascular calcification, that it drives VSMC osteogenic reprogramming, and that genetic or pharmacological interruption of this axis mitigates arterial calcification *in vivo*. Finally, we demonstrate that circulating FNDC1 correlates with tissue calcification and predicts cardiovascular risk, underscoring its translational potential as both a biomarker and therapeutic target.

## MATERIALS AND METHODS

### Calciphylaxis Tissue collection

Human tissue samples were obtained through the Mass General Brigham Calciphylaxis Biorepository and Patient Registry (ClinicalTrials.gov identifier NCT03032835), which prospectively enrolls patients with a clinicopathologically confirmed diagnosis of cutaneous calciphylaxis. Inclusion criteria required participants to be >18 years of age and to have lower extremity involvement. Viable tissue was collected from the margins of active calciphylaxis lesions. Control tissue was obtained from patients undergoing lower extremity amputation for non-calciphylaxis indications. With a two-tailed significance level (α = 0.05), a sample size of 6 calciphylaxis cases and 3 controls provided >85% power to detect differentially expressed genes with a coefficient of variation <0.4. The study was approved by the Institutional Review Board of Massachusetts General Hospital (protocol #2016P002690).

### Human coronary artery tissue procurement

Explanted human coronary artery tissue biospecimens were obtained from heart transplant donors consenting to research studies at Stanford University, as previously described.^17^ Indications for cardiac transplantation included ischemic cardiomyopathy with significant coronary atherosclerosis and non-ischemic cardiomyopathy with coronary arteries exhibiting mild neointimal hyperplasia. Hearts were arrested using cardioplegic solution and transported on ice. Proximal coronary artery segments from the main branches of the left anterior descending, circumflex, or right coronary arteries were dissected. Epicardial and perivascular adipose tissues were trimmed on ice, rinsed in cold phosphate-buffered saline, rapidly frozen in liquid nitrogen, and stored at -80 °C. Non-obstructive human coronary artery tissue biospecimens were also obtained from non-diseased donor hearts rejected for orthotopic heart transplantation, processed following the same protocol as for transplant hearts. Tissues were de-identified, and clinical and histopathologic data were used to classify ischemic and non-ischemic hearts and lesion-containing and non-lesion-containing arteries. Frozen tissue samples were transferred to the University of Virginia under a material transfer agreement and in accordance with Institutional Review Board–approved protocols.

### RNA sequencing and functional enrichment analysis

Differential gene expression analysis was performed using DESeq2, comparing patients with atherosclerotic lesions to those without lesions. Filtered raw read counts and corresponding metadata were provided, with age, sex, and RNA integrity numbers (RIN) used as covariates in the model. Significant differentially expressed genes (DEGs) were identified based on a Benjamini-Hochberg false discovery rate (FDR)-adjusted p-value of <0.05 and a |log2(fold change)| > 1.0. Pathway enrichment analysis was conducted by mapping the DEGs to the Kyoto Encyclopedia of Genes and Genomes (KEGG) database to identify relevant biological pathways. To further explore the functional relationships between the identified DEGs, protein-protein interaction (PPI) network analysis was performed using the STRING database (interaction score > 0.4). The largest subnetwork was selected for functional integration, with KEGG pathways as the reference for enrichment analysis. Network visualization was carried out using Cytoscape v. 3.9.1.

### Hematoxylin and Eosin (H&E) and von Kossa Staining

The slides were deparaffined and hydrated before applying the Harris Hematoxylin solution (Sigma Aldrich HHS 32) for one minute, followed by washing in distilled water for 2 minutes. Eosin was applied for 1-2 minutes as a counterstain. For the von Kossa stain, the deparaffined and hydrated slides were covered with 1% silver nitrate and exposed to bright light for 30 minutes. After washing in distilled water twice for 3 minutes each, 2.5% sodium thiosulphate was applied for 5 minutes. After washing, nuclear fast red was applied for 5 minutes as a counterstain. The slides were then dehydrated and mounted on a coverslip.

### Immunofluorescence microscopy

Immunofluorescence analysis was performed to localize protein expression of DEGs in calciphylaxis tissue and to compare their expression with α-smooth muscle actin (α-SMA), a marker of arterial smooth muscle cells. Paraffin-embedded sections of calciphylaxis tissue were stained according to published protocols.^18^ For immunofluorescent staining of human coronary arteries, fresh frozen cryosections were air-dried at room temperature, fixed with 4% paraformaldehyde for 5 minutes, and washed with PBS. After permeabilization with 0.1% Triton-X, sections were blocked in 10% donkey serum and incubated with primary and secondary antibodies. Primary antibodies consisted of those targeting FNDC1 (PA556603), KYNU (SC-390360), KCNJ15 (PA553474), THBS1 (SC-393504), HIF1α (SC-53546), IFI-16 (SC-8023), NFIL3 (11773-1-AP), RAB31 (H00011031-M03), and DEPP1 (PA5-58122). Protein expression in calciphylaxis tissue was visualized using fluorescence microscopy, and quantification was performed using ImageJ2 (v. 2.9.0) with separate quantification for each arteriole. The Shapiro-Wilk test was applied to assess normality, and differences in protein expression between control and calciphylaxis tissues were analyzed using either a Student’s t-test or Mann Whitney U test, as appropriate. Statistical analyses were performed using R version 4.4.1.

### SiRNA-mediated knockdown and adenovirus-mediated overexpression of FNDC1 in vitro

Human aortic smooth muscle cells (HVSMC, Cell Applications, Cat # 354-05a) were transfected with siRNA targeting FNDC1 or a scrambled control siRNA (siCTRL) purchased from Dharmacon (SMARTpool, Horizon Discovery). Transfections were carried out using DharmaFECT Lipofectamine Reagent (Horizon Discovery) according to the manufacturer’s instructions. For overexpression experiments, recombinant adenoviruses encoding human FNDC1 (with a Green Fluorescent Protein (GFP) sequence separated by an internal ribosomal entry site to assess transduction efficiency), or eGFP alone, were obtained from Vector Biolabs. HVSMCs were transduced with these adenoviral vectors in either normal or osteogenic media for 24 to 48 hours. Gene expression analysis via qPCR and protein expression via western blot were conducted 6 to 8 days post-transduction.

### Calcification, Proliferation, and Migration Assays of HVMSCs

To induce calcification, HVSMCs were cultured in DMEM supplemented with 10% fetal calf serum, 10 mM β-glycerophosphate, 50 μg/mL L-ascorbic acid, and 10 nM dexamethasone, as previously described.^19^ After 21 days of culture, cells were fixed in 10% formalin and stained with a 2% Alizarin Red solution (pH 4.1-4.3) for 2-5 minutes to assess calcification. Cell proliferation was evaluated using an MTT (3-(4,5-dimethylthiazol-2-yl)-2,5-diphenyltetrazolium bromide) assay, as previously described.^20^ HVSMCs were cultured in either osteogenic or normal media in the presence of siRNA, and cell viability was measured after a defined period.

Increased VSMC migration is a hallmark of the osteogenic phenotypic switch. To assess migration, HVSMCs were transfected with siRNA and cultured in normal or osteogenic media for 24 hours. Cells were then plated into a 3-well silicone insert with two defined cell-free gaps (Ibidi, Cat # 80366) and cultured overnight until confluent. The inserts were subsequently removed, and cell migration into the gaps was monitored over time. Migration was quantified by measuring the extent of cell movement into the cell-free areas using ImageJ software.

### Real time polymerase chain reaction and western blot

Total RNA was extracted from HVSMCs following specified treatments using the phenol/guanidinium method. cDNA synthesis was performed using the High-Capacity cDNA Reverse Transcription Kit (Applied Biosystems, Foster City, CA, USA). Quantitative PCR (qPCR) was conducted to assess relative mRNA expression, with normalization to 18S rRNA. TaqMan Gene Expression Assays (Thermo Fisher Scientific) were utilized for gene-specific amplification. For western blot analysis, HVSMCs were lysed in RIPA buffer containing protease and phosphatase inhibitors (Sigma). Protein concentrations were determined, and 20 μg of protein per sample was mixed with Laemmli loading buffer and resolved on an SDS-PAGE gel. The proteins were transferred to polyvinylidene fluoride (PVDF) membranes and incubated with primary antibodies: FNDC1 (St. John’s Laboratory, STJ195139), RUNX2 (Abcam, EPR14334), ALPL (ProteinTech, 82915-2-RR), p-AKT (Ser473) (Cell Signaling, 9271), AKT (Cell Signaling, 5569), NAMPT (Invitrogen, 362616), NMNAT (Life Technologies, 11399-1-AP), NRK (Invitrogen, PIPA5100912), and GAPDH (Cell Signaling, 2118). After incubation with secondary antibody (fluorescent-dye labeled anti-rabbit IgG IRDye 800CW, LI-COR, Lincoln, NE), protein bands were visualized using the LI-COR Odyssey detection system (LI-COR, Lincoln, NE).

To investigate the mechanistic role of FNDC1, AKT, and NAD biosynthesis, AKT Inhibitor IV (Sigma-Aldrich, 124011), FK866 (MedChemExpress, HY-50876), and β-NMN (MedChemExpress, HY-F0004) were used.

### NAD/NADH Assay

A bioluminescent assay (NAD/NADH-Glo Assay, Promega) was used to quantify total NAD as well as the NAD^+^/NADH ratio per manufacturer’s instructions.

### Mitochondrial Respiration and Glycolytic Stress Assay

Mitochondrial respiration and glycolytic function were assessed using the Seahorse XFe96 extracellular flux analyzer (Agilent Technologies, 102416). HVSMCs, previously transduced with either adCTRL or adFNDC1, were seeded at a density of 2 × 10⁴ cells per well in Seahorse XF96 cell culture microplates (Agilent, 101085–004) one day prior to the assay. Cells were plated in bicarbonate-free Dulbecco’s Modified Eagle Medium (DMEM; pH 7.4) supplemented with 4 mM glutamine, 1 mM pyruvate, and 10 mM glucose and incubated overnight at 37°C in a non-CO₂ incubator.

On the day of the assay, sensor cartridges were hydrated and loaded with mitochondrial stress test compounds into the designated injection ports: 1.5 μM oligomycin (injection A), 1 μM FCCP (carbonyl cyanide-4-(trifluoromethoxy)phenylhydrazone; injection B), and 0.5 μM rotenone/antimycin A (injection C), in accordance with the manufacturer’s instructions. Real-time measurements of oxygen consumption rate (OCR) and extracellular acidification rate (ECAR) were obtained using Wave software, and data were analyzed to evaluate mitochondrial and glycolytic function.

### Co-immunoprecipitation Assay (Co-IP)

HVSMCs were cultured in normal media for 3–5 days. Following PBS washes, cells were detached using a cell scraper, and the cell pellet was prepared and lysed as previously described.^21^ For immunoprecipitation, primary antibodies for FNDC1 (Invitrogen PA5-56603) and NAMPT (Invitrogen PA1-1045) were coupled to beads using the Dynabeads Antibody Coupling Kit (Invitrogen 14311D), following the manufacturer’s instructions. The antibody-coupled beads were incubated with the protein lysate overnight at 4°C. After incubation, the beads were washed with appropriate buffers and protein complexes were eluted. The eluted samples were concentrated using centrifugal filters (30 kDa MWCO, Millipore UFC9030) prior to western blot analysis.

### Complex Structure Prediction by Alphafold 3

The protein sequences for FNDC1 (UniProt ID: Q4ZHG4) and NAMPT (UniProt ID: P43490) were retrieved from the UniProt database (https://www.uniprot.org).^22^ The three-dimensional structure of the FNDC1 and NAMPT complex was predicted using AlphaFold 3 (http://alphafoldserver.com) at the various molar ratios outlined in Table S3.^23^ Structural visualization of the complex was performed using PyMOL.^24^

### Animal care

All animal experiments described in this study were approved by the Institutional Animal Care and Use Committees of Massachusetts General Hospital and conducted in accordance with the Guide for the Care and Use of Laboratory Animals of the National Institutes of Health. Fndc1-deficient mice were generated by the deletion of exon 4 using two guide RNAs (Cyagen Biosciences). To investigate the role of FNDC1 in vascular calcification, Mgp^+/-^ mice were crossed with Fndc1^+/−^ and Fndc1^−/−^ mice to ultimately produce Mgp^-/-^ Fndc1^-/-^ and Mgp^-/-^ Fndc1^+/−^ offspring. Animals were maintained on a standard diet. Aortic calcification assessment and survival studies were conducted as detailed below.

To induce vascular calcification, mice were administered vitamin D3 (500,000 IU/kg/day, Sigma-Aldrich) in ethanol mixed with Cremophor (Sigma-Aldrich) and dextrose solution (0.04g/mL) as previously described.^25^ Mice received daily intraperitoneal injections of the vitamin D solution for 3 days and were then maintained on a standard diet for 7 days to allow for the induction of vascular calcification.

### Aortic calcification assessment

Aortic tissue was harvested at 21 days of age for Mgp^-/-^ mice and 10 days post-vitamin D injection for adult mice (6–8 weeks of age). One day prior to euthanasia, mice received a tail vein injection of the fluorescent imaging probe OsteoSense-680 (PerkinElmer, 2 μL/g), as previously described.^26^ Aortas were carefully dissected, excised, and imaged using the Odyssey Imaging System (LI-COR Biotechnology, Lincoln, NE). Aortic fluorescence was normalized to age-matched wild-type mouse aortas, which were used as negative controls. Following imaging, aortas were embedded and cryopreserved in optimal cutting-temperature (OCT) medium (Sakura Tissue-Tek, Zoeterwoude, Netherlands). Serial 5 μm sections were prepared for Alizarin Red staining. The percentage of the vessel area stained with Alizarin Red was quantified using ImageJ.

### Coronary Artery Calcification Analysis in MESA Cohort

To evaluate the association between circulating FNDC1 levels and CAC, we analyzed data from the fifth examination visit of the Multi-Ethnic Study of Atherosclerosis (MESA). FNDC1 protein levels were quantified using Olink’s proximity extension assay and expressed in Normalized Protein eXpression (NPX) units. CAC was assessed using non-contrast cardiac computed tomography and quantified as Agatston scores, as previously described.^27^ Estimated glomerular filtration rate (eGFR) was calculated from serum creatinine concentrations using the CKD-EPI 2021 equation, and chronic kidney disease (CKD) was staged as follows: stage 1 (eGFR > 90), stage 2 (eGFR 60–89), stage 3 (eGFR 30–59), stage 4 (eGFR 15–29), and stage 5 (eGFR < 15 mL/min/1.73 m²). Diabetes mellitus (DM) was defined as impaired fasting glucose, physician diagnosis, or use of anti-diabetic medication. Hypertension was defined by meeting the Joint National Committee (JNC VI) criteria or current use of antihypertensive medication. A multivariable linear regression model was constructed in R (version 4.4.0) to assess the relationship between FNDC1 and CAC, using the following formula:

log₁₀(CAC + 1) ∼ FNDC1 + Age + Sex + CKD + DM + HTN

Covariates were selected based on known associations with vascular calcification.

### Myocardial Infarction Risk Modeling in the UKBiobank

To assess the association between circulating FNDC1 levels and risk of myocardial infarction (MI), we analyzed baseline data from the initial assessment visit of the UK Biobank cohort. FNDC1 protein levels were quantified using Olink’s proximity extension assay and reported in Normalized Protein eXpression (NPX) units. A Cox proportional hazards regression model was constructed to evaluate time to incident MI, with serial inclusion of clinical covariates. The final model included the following variables:

MI ∼ FNDC1 + Age + Sex + CKD + DM + HTN + Body Mass Index (BMI) + Statin Use + Smoking Status

CKD stage was calculated from serum creatinine using the CKD-EPI 2021 equation and classified as previously described. Diagnoses of diabetes and hypertension were included if established prior to or within three months of the baseline visit. Medication use (statins) and smoking status were based on self-reported data at enrollment.

### Statistical analysis

Statistical analyses were performed using R version 4.4.1 and GraphPad Prism 10.3 (GraphPad Software, La Jolla, CA). The Shapiro-Wilk test was employed to assess the normality of variables. Data are presented as mean ± standard error of the mean (s.e.m.) for continuous variables and as n (%) for categorical variables, unless otherwise indicated. Statistical significance was determined using paired two-tailed Student’s t-tests or two-tailed one-way analysis of variance (ANOVA) followed by Sidak post-hoc testing. All in vitro experiments were conducted in triplicate. Kaplan-Meier survival curves with log-rank comparisons were used to evaluate survival differences between Mgp^-/-^Fndc1^-/-^, Mgp^-/-^Fndc1^+/-^, and Mgp^-/-^Fndc1^+/+^ mouse models.

## RESULTS

### Transcriptomic profiling of calciphylaxis reveals dysregulation of calcification, inflammation, and thrombosis pathways

To elucidate molecular mechanisms underlying aberrant calcification in calciphylaxis, we conducted RNA sequencing of lower extremity skin and soft tissue specimens obtained from patients with and without calciphylaxis undergoing limb amputation (Table S1). In individuals with calciphylaxis, samples were collected from macroscopically viable non-ulcerated regions adjacent to areas of tissue necrosis. Differential gene expression analysis identified 543 significantly upregulated and 240 downregulated transcripts in calciphylaxis tissue relative to controls (Fig. 1A). The most prominently upregulated genes included classical pro-calcific mediators *RUNX2*, a transcription factor essential for osteogenic reprogramming of vascular smooth muscle cells and skeletal development (fold-change 5.69; Q = 8.61 × 10⁻⁴; Table S2)^19,28,29^ and alkaline phosphatase (*ALPL*), an enzyme that hydrolyzes pyrophosphate-a potent endogenous inhibitor of mineralization.(fold-change 25.85; Q = 5.28 × 10⁻⁵; Table S2).^30–32^ Gene set enrichment analysis highlighted robust activation of canonical calcification pathways, including osteoclast differentiation, Wnt signaling, and calcium-phosphate metabolism (Fig. 1B).

**Figure 1.**
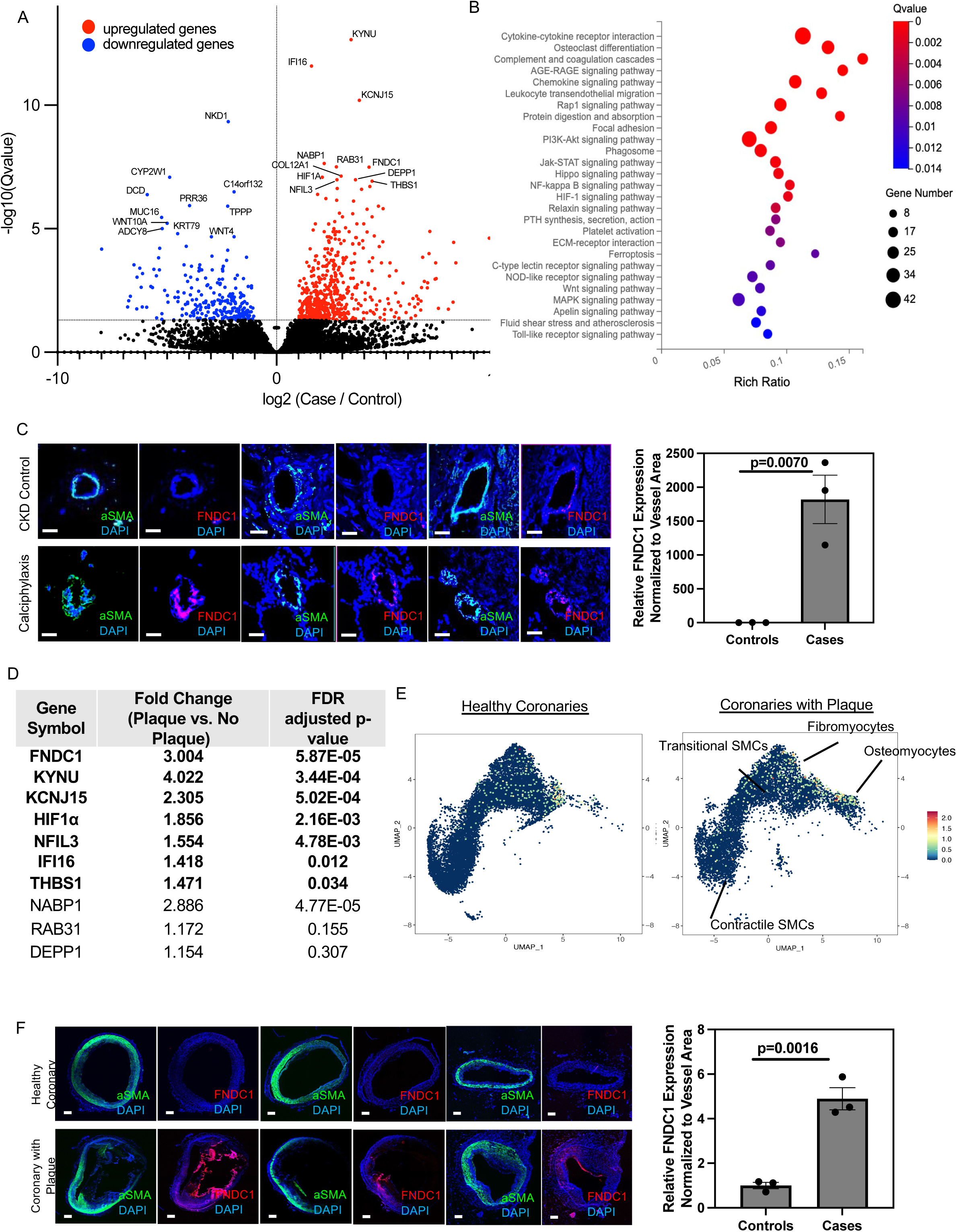
FNDC1 is a top upregulated gene in calciphylaxis and coronary artery disease and is enriched in vascular smooth muscle cells (VSMCs). **A**. Volcano plot of differentially expressed genes (DEGs) comparing calciphylaxis to control tissue. A total of 543 genes were significantly upregulated (red) and 240 genes were significantly downregulated (blue) based on thresholds of FDR-adjusted *p* ≤ 0.05 and |log₂(fold change)| > 1. Select top DEGs are labeled. **B.** Enriched signaling pathways in calciphylaxis tissue (FDR-adjusted p-value [Q-value] ≤ 0.05). Bubble size denotes the number of genes involved in each pathway; color scale reflects statistical significance. Rich ratio indicates the proportion of genes altered within each pathway (see Table S3 for full pathway statistics). **C**. Representative immunofluorescence images of FNDC1 (red), *α*-smooth muscle actin (*α*-SMA, green), and DAPI (blue) in arterioles from calciphylaxis biopsies and matched control tissue. FNDC1 protein levels were significantly increased in calciphylaxis (*p* = 0.0070). Scale bar, 200 μm. **D.** Expression of top upregulated genes from calciphylaxis assessed in coronary arteries with and without atherosclerotic plaque. Genes are ranked by statistical significance. **E**. UMAP visualization of FNDC1 expression across VSMC subpopulations, demonstrating increased expression in plaque-containing coronary arteries (bottom) relative to plaque-free arteries (top). FNDC1 expression increased with phenotypic modulation of VSMCs from contractile to fibromyocyte and osteogenic-like states. **F.** Representative immunofluorescence images of FNDC1 (red), α-SMA (green), and DAPI (blue) in coronary arteries with and without plaque. FNDC1 protein expression was significantly higher in plaque-bearing coronaries (*p* = 0.0016). Scale bar, 200 μm.

Pathway-level analysis further demonstrated significant enrichment of inflammatory and thrombotic signaling cascades, consistent with established histopathologic features of calciphylaxis and overlapping with molecular signatures observed in athero-calcific lesions of large-vessel disease.^8,33,34^ Pro-inflammatory mediators such as *IL1β* and *IL6* were markedly upregulated in calciphylaxis tissue (fold-change 36, Q = 2.43 × 10⁻⁴ and fold-change 21, Q = 4.98 × 10⁻⁴, respectively), supporting their mechanistic role in vascular calcification and immune-mediated tissue injury.^35–38^ We also observed elevated expression of *SERPINE1* (encoding plasminogen activator inhibitor 1), as well as classical complement components *C1QA* and *C1QB*, implicating complement-coagulation axis activation in disease pathogenesis (Table S3).^39^ Additional dysregulated pathways included those governing autophagy (phagosome) and redox balance (AGE-RAGE signaling). Network analysis of upregulated transcripts revealed a core module of 479 genes and 24 significantly enriched pathways, encompassing inflammation, mineralization, thrombosis, autophagy, and cell-cell interactions. Among the top differentially expressed genes (DEGs), eight of the nine most upregulated-including *FNDC1* (fibronectin type III domain-containing 1)-occupied central positions within this network, suggesting key roles in orchestrating the transcriptomic landscape of calciphylaxis-associated vasculopathy.

### Vascular FNDC1 expression is upregulated in human calciphylaxis and coronary artery disease

To interrogate the role of top upregulated genes in calciphylaxis pathogenesis, we evaluated their protein expression and spatial localization relative to alpha–smooth muscle actin (α-SMA), a canonical marker of VSMCs, using immunofluorescence in an independent patient cohort. Histopathologic analysis on hematoxylin and eosin (H&E) and von Kossa staining revealed extensive medial calcium deposition in small-to medium-sized subcutaneous vessels in calciphylaxis tissue, findings that were absent in control tissues (Fig. S1). Among the nine most highly upregulated DEGs, seven displayed concordant increases in protein abundance in calciphylaxis specimens relative to controls, validated across three clinically matched independent replicates (Fig. S2). Several of these proteins were highly enriched within the vasculature of affected tissue, exhibiting expression levels multiple orders of magnitude above baseline. Notably, FNDC1 demonstrated one of the most pronounced increases (p = 0.007), with strong vascular localization (Fig. 1C).

To determine whether the transcriptomic alterations observed in calciphylaxis extend to large-vessel disease, we analyzed bulk RNA sequencing data from 40 human coronary arteries with and without atherosclerotic lesions. All seven validated calciphylaxis DEGs were significantly upregulated in diseased coronary arteries relative to controls (Fig. 1D), suggesting a shared molecular signature between microvascular and macrovascular calcification. Protein-level validation in coronary arteries corroborated these findings. Immunofluorescence analysis of FNDC1, KYNU (kynureninase), and KCNJ15 (potassium inwardly rectifying channel subfamily J member 15) revealed elevated expression of all three proteins in calcified vessels (Fig. S3). Among these, FNDC1 exhibited the strongest signal, localizing both to the medial layer and within atherosclerotic plaques (p = 0.0074; Fig. 1F). Single-cell RNA sequencing of atherosclerotic coronary and carotid arteries further demonstrated dynamic upregulation of FNDC1 during the phenotypic transition of VSMCs from a contractile to an osteogenic state (Fig. 1E), consistent with a role for FNDC1 in promoting VSMC phenotypic modulation in CAD. Together, these results demonstrate marked transcriptional and translational convergence between small- and large-vessel calcific disease, positioning FNDC1 as a key molecular mediator in VSMC-driven vascular calcification across distinct vascular beds.

### FNDC1 mediates human vascular smooth muscle cell calcification, proliferation, and migration

VSMC calcification is driven by a phenotypic transition from a contractile to an osteogenic state, marked by upregulation of bone-associated genes such as *RUNX2* and *ALPL*. This osteogenic phenotype is also characterized by enhanced migratory and proliferative capacity, features that contribute to fibrous cap formation in atherosclerotic plaques.^12^ To investigate the functional role of *FNDC1* in mediating the osteogenic switch, we performed knockdown studies in human vascular smooth muscle cells (HVSMCs) cultured under high-phosphate osteogenic conditions. Silencing *FNDC1* using small interfering RNA (siRNA) significantly reduced calcification, as assessed by Alizarin Red staining at 21 to 28 days post-induction, compared to cells transfected with control siRNA (siCTRL) (p = 0.036; Fig. 2A). Treatment with siFNDC1 effectively blunted the induction of *FNDC1* observed under osteogenic conditions and its knockdown was associated with marked suppression of *RUNX2* and *ALPL* (p ≤ 0.0001 for both; Fig. 2, B and C). Functionally, *FNDC1* knockdown also attenuated the increased proliferation and migration observed in HVSMCs cultured in osteogenic media (p < 0.0001 and p = 0.0165, respectively; Fig. 2D and Fig. S4). Comparative silencing of *KYNU* and *KCNJ15*, two additional top DEGs in calciphylaxis, revealed that FNDC1 depletion produced the most robust suppression of calcification, proliferation, and migration (Fig. S5).

**Figure 2.**
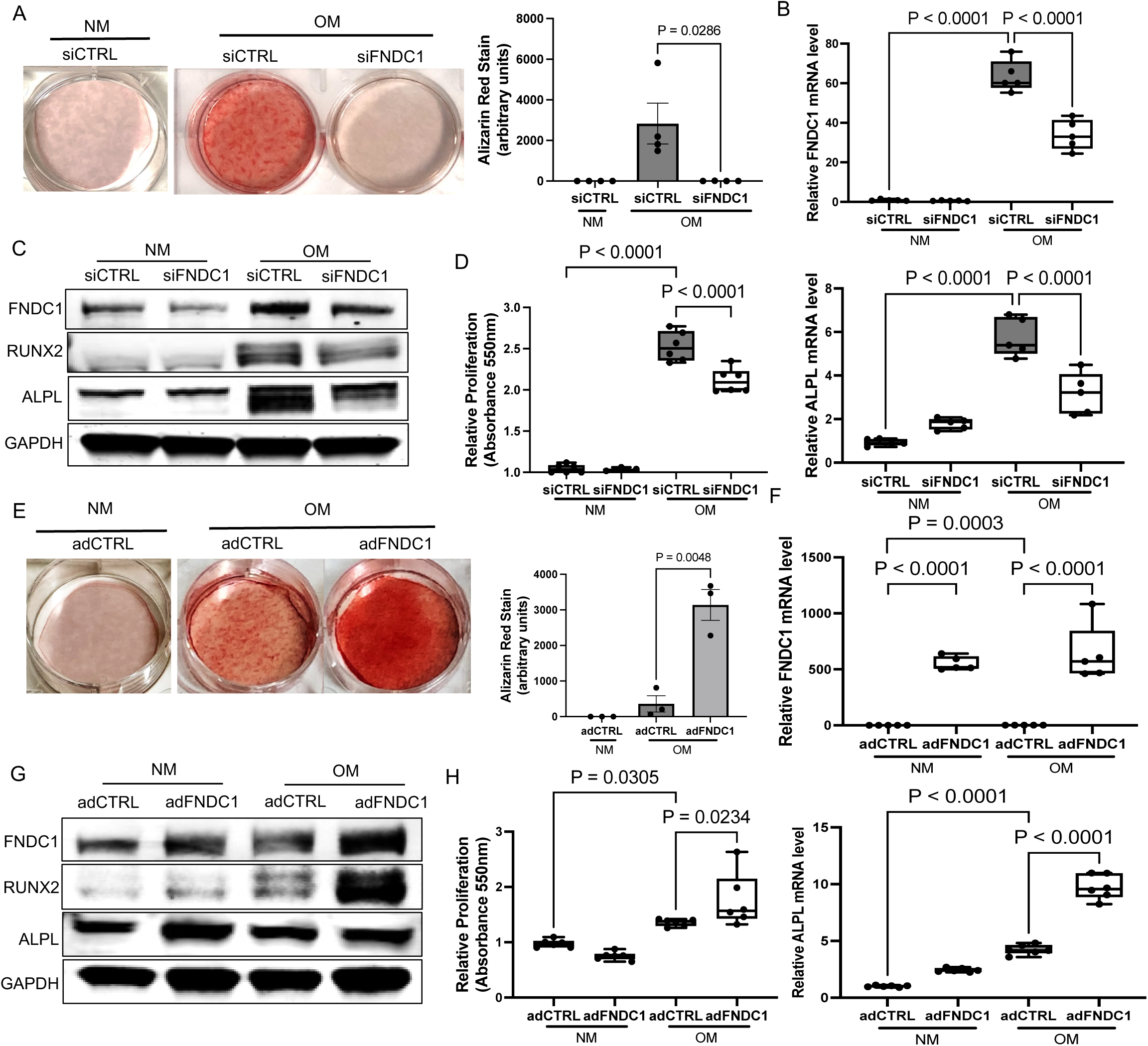
FNDC1 promotes osteogenic phenotypic transition and calcification in human vascular smooth muscle cells (HVSMCs). **A**. Silencing of FNDC1 in HVSMCs cultured in osteogenic media markedly reduced calcium deposition as assessed by Alizarin red staining after 21–24 days. Quantification showed >90% reduction in calcification in siFNDC1-treated cells compared to siCTRL (*p* = 0.0286). **B.** FNDC1 mRNA expression was significantly upregulated in HVSMCs after 3 days of culture in osteogenic media. siFNDC1 treatment reduced FNDC1 mRNA levels by ∼50%, which was accompanied by a significant downregulation of *ALPL* expression (*p* ≤ 0.0001). **C**. Western blot of HVSMCs cultured in normal or osteogenic media following transfection with siCTRL or siFNDC1. FNDC1 knockdown attenuated the osteogenic media-induced upregulation of RUNX2 and ALPL protein expression. **D**. Cell proliferation assessed by MTT assay after 3 days of culture in normal or osteogenic media. siFNDC1 significantly suppressed HVSMC proliferation in osteogenic conditions (*p* < 0.0001). **E.** HVSMCs transduced with adenoviral FNDC1 (adFNDC1) exhibited increased calcification after 10-14 days in osteogenic media compared to adCTRL-transduced cells, as visualized by Alizarin red staining (*p* = 0.0048). **F**. Quantitative PCR revealed significantly increased *ALPL* expression in adFNDC1-transduced HVSMCs cultured in osteogenic media relative to adCTRL (*p* < 0.0001). **G.** Overexpression of FNDC1 increased RUNX2 and ALPL protein levels in HVSMCs, as shown by Western blot. **H.** FNDC1 overexpression enhanced HVSMC proliferation in osteogenic media, as measured by MTT assay (*p* = 0.0234). All statistical analyses were performed with one-way ANOVA with Sidak’s post-hoc comparison test.

To further define the pro-calcific role of FNDC1, we overexpressed *FNDC1* in HVSMCs using an adenoviral vector (adFNDC1). Compared to cells transduced with a control vector (adCTRL), adFNDC1-transduced HVSMCs exhibited significantly accelerated calcification in osteogenic media, with mineral deposition observed after ∼10 days (p = 0.0048; Fig. 2E). This represented a reduction in the time to calcification onset by more than 50%, highlighting the strong calcification-promoting effect of *FNDC1*. *FNDC1* overexpression also led to increased expression of *RUNX2* and *ALPL* under both basal and osteogenic conditions (Fig. 2, F and G), and significantly enhanced proliferative capacity in osteogenic media (p = 0.0234; Fig. 2H). Collectively, these results support a central role for *FNDC1* in osteogenic reprogramming and functional activation of VSMCs, with a coordinated promotion of calcification, proliferation, and migration.

### FNDC1 depletion ameliorates disease in two different mouse models of vascular calcification

To investigate the *in vivo* role of *FNDC1* in vascular calcification, we generated an Fndc1 knockout mouse model (Fndc1^-/-^) using CRISPR-Cas9–mediated deletion of exon 4 (Fig. S6). *Fndc1* knockout mice were evaluated in two independent models of vascular calcification: the matrix Gla protein–deficient (*Mgp^-/-^*) mouse model and the vitamin D-induced mouse model.^14,25,40^ *Mgp^-/-^* mice develop widespread spontaneous arterial calcification by postnatal day 14 and succumb to aortic rupture by 5 to 6 weeks of age.^14,40^ Quantitative PCR revealed significantly elevated *Fndc1* expression in the aortas of *Mgp^-/-^* mice compared to wild-type (WT) littermates at both 14 days (p = 0.0022) and 28 days of age (p = 0.0095; Fig. 3A). Immunofluorescence staining confirmed increased expression of FNDC1 in *Mgp^-/-^* aortas relative to age-matched WT controls (p = 0.0062 and p = 0.0036, respectively; Fig. 3B).

**Figure 3.**
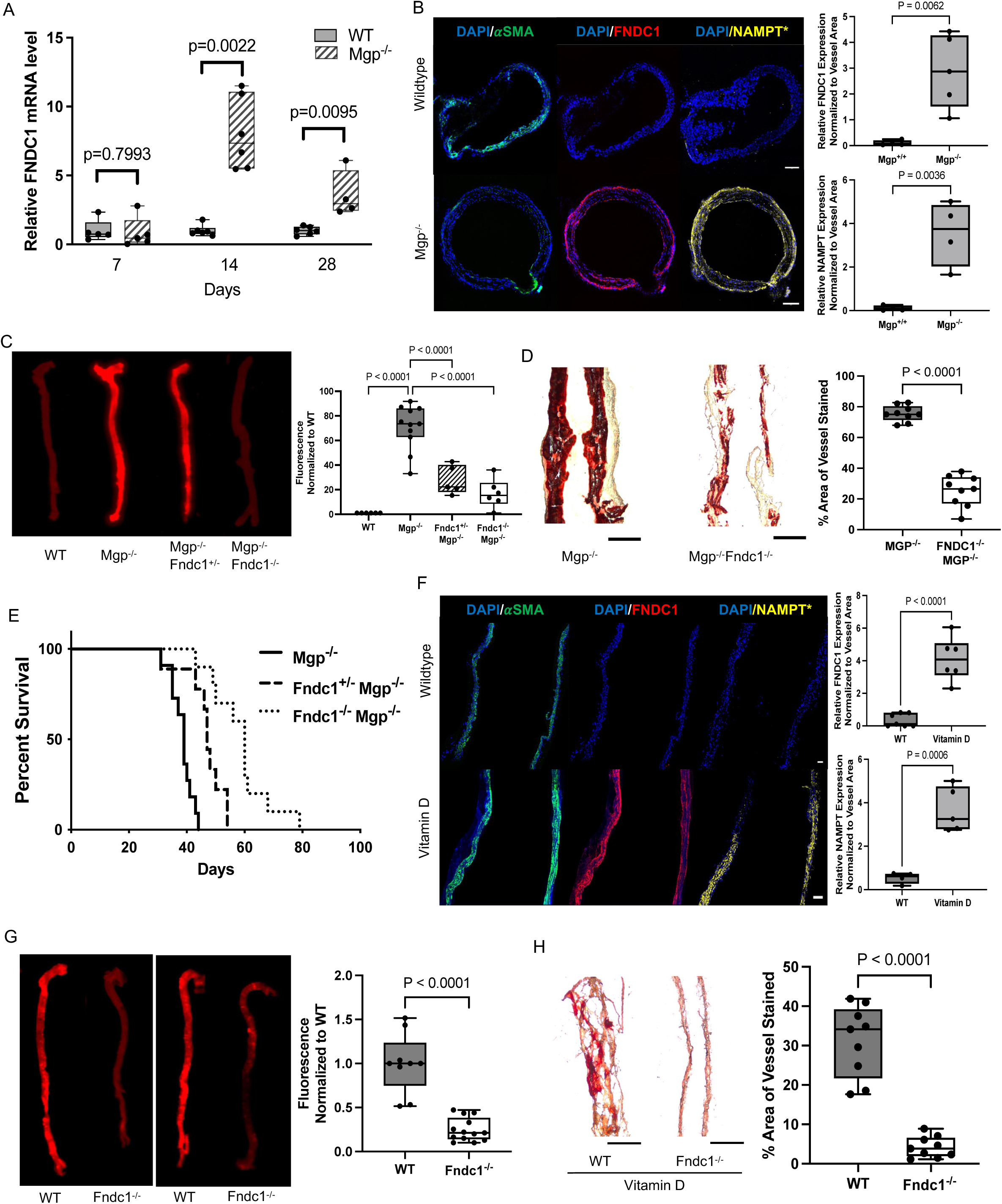
Genetic depletion of FNDC1 reduces vascular calcification and improves survival in murine models of arterial calcification. **A**. Quantitative PCR analysis of aortic tissue from *Mgp^-/-^*mice demonstrated increased *Fndc1* mRNA expression at postnatal days 14 and 28 compared to age-matched wild-type (WT) controls. **B.** Immunofluorescence images of aortic sections showed increased FNDC1 (red) and NAMPT (yellow) expression in *Mgp^-/-^* mice relative to WT mice. *Indicates serial section (5 μm) stained for NAMPT. **C.** Near-infrared fluorescence imaging (NIRF) revealed significantly reduced vascular calcification in *Mgp^-/-^ Fndc1^+/-^* and *Mgp^-/-^ Fndc1^-/-^* mice compared to *Mgp^-/-^Fndc1^+/+^* controls (*p* < 0.0001 for both comparisons). **D**. Alizarin red staining of aortic tissue confirmed decreased mineral deposition in *Mgp^-/-^ Fndc1^-/-^* mice relative to *Mgp^-/-^Fndc^+/+^* littermates (*p* < 0.0001). **E**. Kaplan–Meier survival analysis showed improved survival in *Mgp*^-/-^ *Fndc1*^-/-^ mice (*n* = 10, *p* < 0.0001) and *Mgp*^-/-^ *Fndc1^+^*^/-^ mice (*n* = 9, *p* = 0.0004) compared to *Mgp*^-/-^ *Fndc1*^+/+^ controls (*n* = 11, log-rank *p* < 0.0001 and *p* = 0.0004, respectively). **F.** Immunofluorescence staining demonstrated increased FNDC1 (red) and NAMPT (yellow) expression in the aortas of vitamin D-treated WT mice compared to vehicle controls. **G.** NIRF imaging of aortas showed reduced vascular calcification in vitamin D-injected *Fndc1^-/-^* mice compared to WT mice (*p* < 0.0001). **H**. Alizarin red staining corroborated the decrease in vascular calcification in *Fndc1^-/-^* mice following vitamin D administration (*p* < 0.0001). Scale bar, 200 μm. Unless otherwise indicated, all statistical analyses were performed with either the unpaired t-test for two group comparisons or one-way ANOVA with Sidak’s post-hoc comparisons test for more than two groups.

Remarkably, *Mgp^-/-^Fndc1^-/-^* double-knockout mice exhibited >60% reduction in aortic calcification, as quantified by near-infrared fluorescence imaging and alizarin red staining (p < 0.0001 for both; Fig. 3, C and D). This reduction was accompanied by a significant extension in median survival—from 39 days in *Mgp^-/-^* mice to 60 days in *Mgp^-/-^Fndc1^-/-^* littermates (log-rank p = 0.0013; Fig. 3E). *Mgp^-/-^Fndc^+/-^* heterozygous mice demonstrated intermediate reductions in aortic calcification (∼30% reduction; p < 0.0001) and improved median survival (47.5 days; log-rank p = 0.0004; Fig. 3, C–E), indicating a dose-dependent effect of *Fndc1* deletion on vascular mineralization and lifespan. In the vitamin D-induced vascular calcification model, FNDC1 was also significantly upregulated in the aortas of vitamin D–treated mice relative to vehicle-treated WT controls (p < 0.0001 and p = 0.006, respectively; Fig. 3F). Consistent with findings from the *Mgp^-/-^* model, vitamin D-treated *Fndc1^-/-^* mice exhibited >70% reduction in aortic calcification, as assessed by both near-infrared imaging and alizarin red staining (p < 0.0001 for both; Fig. 3, G and H). These findings establish *FNDC1* as a critical regulator of vascular calcification across multiple *in vivo* models.

### FNDC1-dependent VSMC osteogenesis is mediated by AKT and NAMPT

Recent studies have identified FNDC1 as a regulator of oncogenic signaling, including its direct interaction with the G_βγ_ subunit of heterotrimeric G proteins, leading to activation of downstream pathways such as PI3K-AKT that support tumor proliferation and metastasis.^41,42^ As an established key regulator of cellular growth, migration, and survival, the PI3K-AKT pathway is also critical to the osteogenic phenotypic transition of VSMCs. Phosphorylated AKT is a known activator of RUNX2 and multiple intracellular signaling programs linked to calcification and VSMC plasticity.^43,44^ To determine whether *FNDC1* promotes VSMC calcification through PI3K-AKT activation, we assessed AKT phosphorylation in HVSMCs following either *FNDC1* knockdown or overexpression under both normal and osteogenic conditions. Silencing *FNDC1* attenuated AKT phosphorylation under osteogenic conditions, whereas FNDC1 overexpression enhanced AKT phosphorylation in both basal and osteogenic media, indicating that FNDC1 activates AKT signaling independent of phosphate-induced stimuli (Fig. 4A). Pharmacologic inhibition of AKT (AKT inhibitor IV)^45^ significantly reduced *FNDC1*-driven calcification (p < 0.0001; Fig. 4B), an effect that was accompanied by downregulation of RUNX2 and ALPL expression (Fig. 4C). FNDC1-mediated enhancement of HVSMC proliferation in osteogenic media was similarly abrogated by AKT inhibition (Fig. 4D), implicating PI3K-AKT signaling in FNDC1-induced VSMC phenotypic modulation.

**Figure 4.**
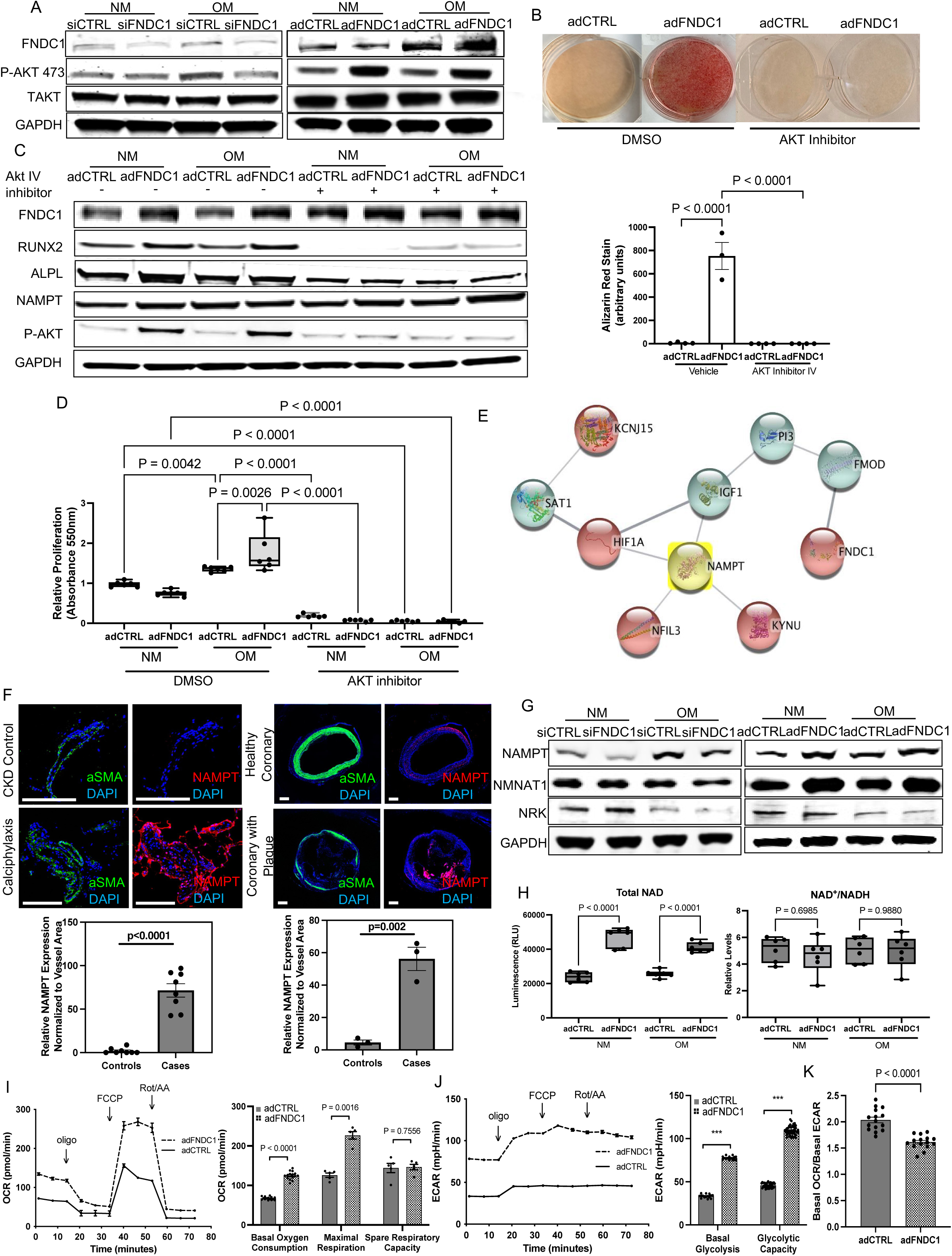
FNDC1 promotes vascular calcification through activation of PI3K–AKT signaling and modulates the metabolic phenotype of HVSMCs through regulation of NAMPT. **A**. Western blot analysis of HVSMCs transfected with siFNDC1 (left) or transduced with adenoviral FNDC1 (adFNDC1; right). FNDC1 knockdown decreased phosphorylation of AKT at serine 473, whereas overexpression of FNDC1 increased phospho-AKT levels relative to controls. **B.** Pharmacologic inhibition of AKT signaling with AKT inhibitor IV (100 nM) significantly reduced adFNDC1-induced calcification in HVSMCs, as assessed by Alizarin red staining (*p* < 0.0001). **C**. AKT inhibitor IV (100 nM) attenuated the upregulation of RUNX2 and ALPL protein levels in HVSMCs induced by adenovirus-mediated expression of FNDC1. However, AKT inhibition did not affect NAMPT expression. **D**. Adenoviral expression of FNDC1 increased HVSMC proliferation in osteogenic media (*p* = 0.0026), while AKT inhibition reduced proliferation in both control and FNDC1-overexpressing cells (*p* < 0.0001 for all comparisons). **E.** Protein-protein interaction network analysis of differentially expressed genes in calciphylaxis demonstrate convergence of 5 of the top upregulated genes in calciphylaxis on NAMPT. **F**. Immunofluorescece analysis demonstrated increased NAMPT expression in arterioles from patients with calciphylaxis and in human coronary arteries with plaque compared to respective controls (*p* < 0.0001 and *p* = 0.002, respectively). **G.** Western blot analysis of HVSMCs transfected with siFNDC1 or transduced with adFNDC1 in the presence or absence of osteogenic media. siFNDC1 reduced NAMPT and NMNAT expression under osteogenic conditions, whereas adFNDC1 upregulated both proteins. There were no significant changes in NRK expression with either knockdown or overexpression of FNDC1. **H.** Quantification of intracellular NAD using the NAD^+^/NADH Glo bioluminescence assay (Promega) revealed increased total NAD content in HVSMCs overexpressing FNDC1 under both normal and osteogenic conditions. No significant differences in the NAD^+^/NADH ratio were observed between adFNDC1- and adCTRL-treated cells. **I**. Mitochondrial respiration was assessed by measuring oxygen consumption rate (OCR) before and after sequential injections of oligomycin (oligo, 1.5 μM), FCCP (1 μM), and rotenone/antimycin A (Rot/AA, 0.5 μM) using the Agilent Seahorse XFe24 Analyzer. Basal respiration (OCR prior to oligomycin) and maximal respiration (OCR following FCCP) were elevated in HVSMCs overexpressing FNDC1 compared to controls. **J.** Glycolytic flux was evaluated by extracellular acidification rate (ECAR) measurements before and after oligomycin treatment. FNDC1 overexpression increased both basal glycolysis and glycolytic capacity in HVSMCs. **K.** The ratio of basal OCR to basal ECAR was reduced in FNDC1-overexpressing cells, indicating a metabolic shift favoring glycolysis over oxidative phosphorylation.

To further delineate the downstream molecular pathways through which *FNDC1* promotes vascular calcification, we performed protein-protein interaction analysis of the top DEGs identified in calciphylaxis tissue. This analysis identified nicotinamide phosphoribosyltransferase (*NAMPT*) as a central node linking 5 of the top 9 upregulated genes, including *FNDC1*, *KYNU*, *KCNJ15*, *HIF1α* (hypoxia inducible factor 1 subunit alpha), and *NFIL3* (nuclear factor interleukin 3) (Fig. 4E). *NAMPT* encodes the rate-limiting enzyme in the nicotinamide adenine dinucleotide (NAD⁺) salvage pathway and catalyzes the formation of β-nicotinamide mononucleotide (β-NMN) from nicotinamide.^46^ In our transcriptomic analysis of calciphylaxis tissue, *NAMPT* was markedly upregulated (fold-change = 11.14, Q-value = 2.5 × 10⁻⁶; Table S2), and immunofluorescence staining demonstrated elevated NAMPT protein levels in arterioles from patients with calciphylaxis and in coronary arteries with atherosclerotic plaque compared to healthy controls (Fig. 4F). Given the prominent positioning of *NAMPT* within the *FNDC1*-centered calciphylaxis network and its marked upregulation in diseased tissue, we next investigated whether *FNDC1* modulates VSMC calcification, at least in part, through *NAMPT*-dependent mechanisms.

We first assessed NAD⁺ biosynthesis pathway components in HVSMCs. *NAMPT* protein expression was increased in HVSMCs cultured in osteogenic media, an effect that was significantly attenuated by *FNDC1* knockdown (Fig. 4G). Expression of nicotinamide mononucleotide adenylyltransferase (NMNAT), an enzyme immediately downstream of *NAMPT* that catalyzes the conversion of NMN to NAD⁺, was similarly reduced following *FNDC1* silencing. In contrast, expression of nicotinamide riboside kinase (NRK), which participates in an alternative NAD⁺ salvage pathway utilizing nicotinamide riboside (NR) rather than nicotinamide, was upregulated upon *FNDC1* knockdown despite being generally suppressed under osteogenic conditions (Fig. 4G). Conversely, *FNDC1* overexpression induced robust increase in both *NAMPT* and *NMNAT* expression, with no significant change in *NRK* (Fig. 4G).

To evaluate whether *FNDC1*-mediated VSMC calcification via *NAMPT* involves alterations in cellular redox status, we quantified total NAD(H) and oxidized NAD⁺ levels in HVSMCs cultured in both normal and osteogenic media. Overexpression of *FNDC1* led to a significant increase in total intracellular NAD(H), without affecting the NAD⁺/NADH ratio (Fig. 4H). To correlate elevated intracellular NAD levels with overall VSMC cellular metabolism, we evaluated the real-time extracellular metabolic flux profile of HVSMCs overexpressing FNDC1 using the Agilent Seahorse XFe96 Analyzer. FNDC1 overexpression increased both basal and maximal oxygen consumption rates (OCR), indicating enhanced mitochondrial respiration (Fig. 4I). In parallel, extracellular acidification rate (ECAR) measurements demonstrated increased basal glycolysis and glycolytic capacity in adFNDC1-transduced cells, consistent with elevated glycolytic flux (Fig. 4J). Interestingly, although there was a rise in both OCR and ECAR with adFNDC1, the OCR/ECAR ratio was significantly reduced, suggesting that FNDC1 mediates a metabolic shift in favor of glycolysis over oxidative phosphorylation (Fig. 4K).

FK866 is a selective molecular inhibitor of NAMPT that depletes the cytoplasmic NAD pool and that has been evaluated in clinical trials for cancer. ^47–49^ In our study, pharmacologic inhibition of *NAMPT* with FK866 significantly reduced *FNDC1*-induced calcification (p < 0.0001; Fig. 5A), and this reduction was accompanied by decreased protein expression of the osteogenic markers RUNX2 and ALPL (Fig. 5B). Together, these findings support a role for *NAMPT*-dependent NAD⁺ biosynthesis in mediating *FNDC1*-driven VSMC calcification. Notably, inhibition of AKT signaling did not alter NAMPT expression in FNDC1-overexpressing cells, nor did NAMPT inhibition affect AKT phosphorylation (Figs. 4C and 5B), indicating that PI3K-AKT and NAD⁺ biosynthesis function as parallel, non-redundant pathways downstream of FNDC1.

**Figure 5.**
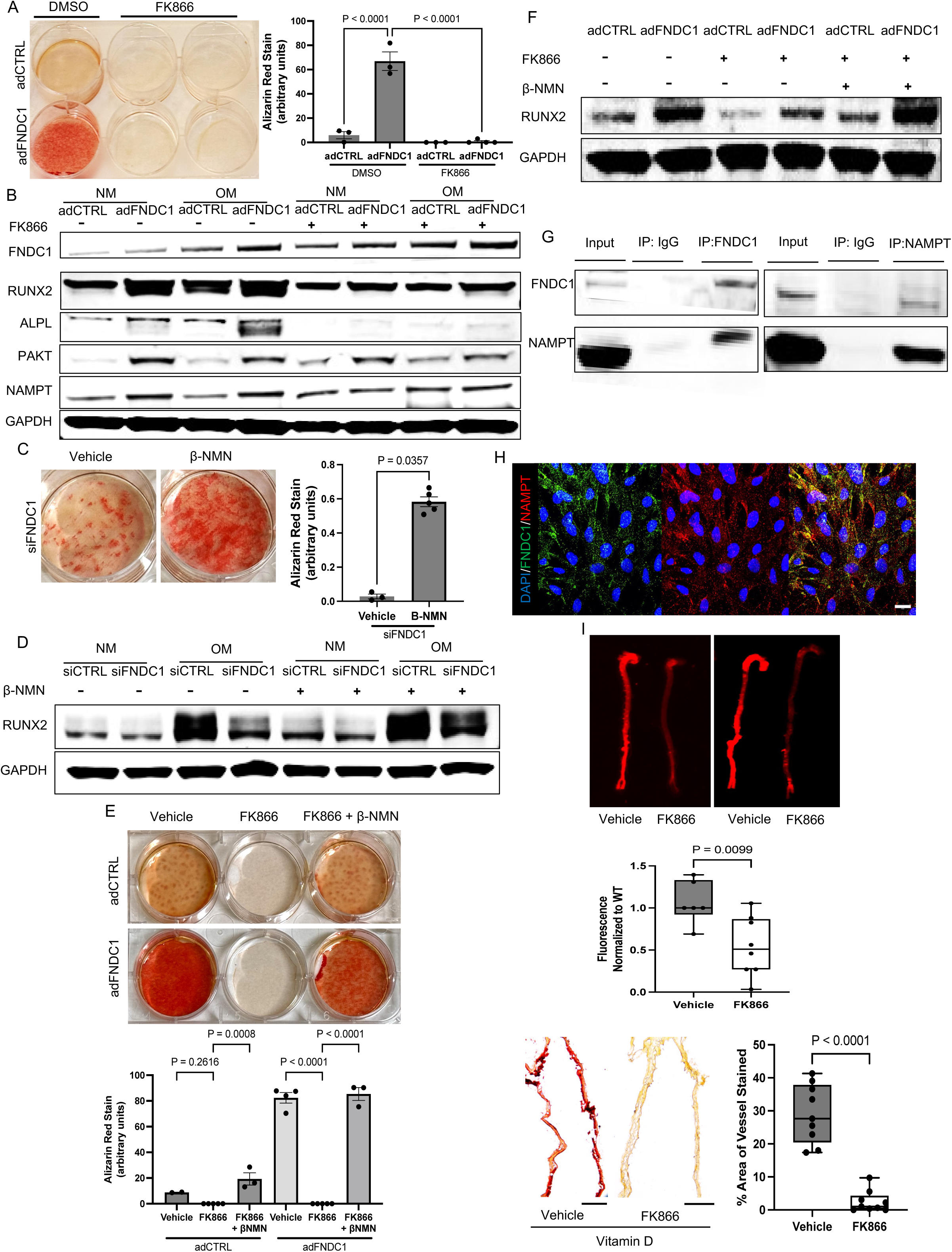
FNDC1 promotes RUNX2-dependent HVSMC calcification through NAMPT. **A.** Pharmacologic inhibition of NAMPT with FK866 (10nM) attenuated calcification induced by adenoviral FNDC1 overexpression in HVSMCs (*p* < 0.0001). **B.** Western blot analysis showed reduced expression of RUNX2 and ALPL in adFNDC1-transduced HVSMCs following FK866 treatment compared to vehicle treatment. **C.** Supplementation with βNMN restored calcification in HVSMCs transfected with siFNDC1 compared to vehicle-treated controls (*p* = 0.0357). **D.** βNMN supplementation in siFNDC1-transfected HVSMCs led to upregulation of RUNX2 expression, consistent with increased calcification. **E.** FK866 treatment reduced FNDC1-induced calcification, while co-treatment with βNMN restored calcification to levels observed in vehicle-treated cells. **F.** Reflecting the changes in calcification, RUNX2 expression was decreased by FK866 treatment in FNDC1-overexpressing HVSMCs and restored by βNMN supplementation. **G.** Co-immunoprecipitation of total protein from HVSMCs cultured in normal media demonstrated a physical interaction between FNDC1 and NAMPT. NAMPT was detected in FNDC1 immunoprecipitates and vice versa. **H.** Immunofluorescence microscopy of HVSMCs cultured in normal media showed colocalization of FNDC1 (green) and NAMPT (red), quantified by Mander’s coefficients (FNDC1 = 0.192). **I.** In wild-type mice subjected to vitamin D-induced calcification, FK866 administration (20 mg/kg) was performed intraperitoneally every 3 days until the time of harvest 7 days after the last vitamin D injection. FK866 treatment reduced aortic calcification, as assessed by near-infrared fluorescence imaging (upper panel) and on Alizarin red staining (lower panel).

Next, to test whether NAD⁺ supplementation is sufficient to restore calcification in the context of FNDC1-deficiency, we treated *siFNDC1*-transfected HVSMCs cultured in osteogenic media with β-NMN, the enzymatic product of *NAMPT*. β-NMN significantly increased calcification (p = 0.0357; Fig. 5C) and restored RUNX2 expression in the setting of *FNDC1* knockdown (Fig. 5D). Similarly, β-NMN reversed the reduction in calcification induced by FK866 in *FNDC1*-overexpressing HVSMCs, restoring both calcification and RUNX2 protein levels to control values (Fig. 5E-F). These findings establish NAMPT-dependent NAD⁺ biosynthesis as a key effector of FNDC1-driven osteogenic programming in VSMCs.

To investigate the structural basis for a potential FNDC1-NAMPT interaction, we employed AlphaFold 3 to predict the three-dimensional structure of an FNDC1-NAMPT complex at various molar ratios (Table S4). While a complete FNDC1 structure could not be resolved, modeling of the N-terminal region (residues 40–470) identified a high-confidence interaction between FNDC1(360–454) and dimeric NAMPT (ipTM = 0.83; Table S4, Fig. S7A). Despite low sequence homology among the four N-terminal domains of FNDC1, they were structurally conserved (Fig. S7B–C), with domain-specific modeling confirming FNDC1(360– 454) as the most likely NAMPT-interacting segment (Table S5, Fig. S7D).

Co-immunoprecipitation assays in HVSMCs confirmed endogenous physical interaction between FNDC1 and NAMPT (Fig. 5G). Immunofluorescence analysis further demonstrated subcellular colocalization under basal conditions, as quantified by Mander’s colocalization coefficients (FNDC1 = 0.192; NAMPT = 0.031; Fig. 5H), supporting a direct intracellular interaction.

*In vivo*, NAMPT protein expression was elevated in aortas of both *Mgp^-/-^* and vitamin D– treated mice, mirroring FNDC1 expression patterns (Fig. 5I). To evaluate the functional impact of NAMPT inhibition *in vivo*, *Mgp^-/-^* mice were treated with FK866 (20 mg/kg every 3 days) beginning at postnatal day 5. FK866 treatment reduced aortic calcification by ∼50% relative to vehicle-treated controls, as determined by near-infrared fluorescence imaging (p = 0.0099) and histological analysis (p < 0.0001; Fig. 5I). These findings demonstrate that *NAMPT* is a critical downstream effector of *FNDC1* and that pharmacologic targeting of NAD⁺ biosynthesis attenuates vascular calcification *in vivo*.

### Circulating FNDC1 is a biomarker of calciphylaxis and coronary artery calcification

Given the high expression of FNDC1 in calciphylaxis vasculature and diseased coronary vessels as well as its functional role in mediating VSMC calcification both *in vitro* and *in vivo*, we next investigated the potential of FNDC1 to serve as a circulating biomarker of vascular calcification. Using plasma samples from 31 patients with calciphylaxis and 31 chronic kidney disease (CKD)-matched control patients, we observed significantly elevated circulating FNDC1 levels in the calciphylaxis cohort (mean ± SEM 63.9 ± 11.1 pg/mL) compared to the control group (30.4 ± 5.7 pg/mL; Mann-Whitney p < 0.0001, Fig. 6), as measured by a competitive ELISA (Abbexa, Cambridge, UK). Demographic and clinical characteristics, including age, sex, and comorbidities such as hypertension, hyperlipidemia, coronary artery disease (CAD), and diabetes mellitus, were comparable between the two groups.

**Figure 6.**
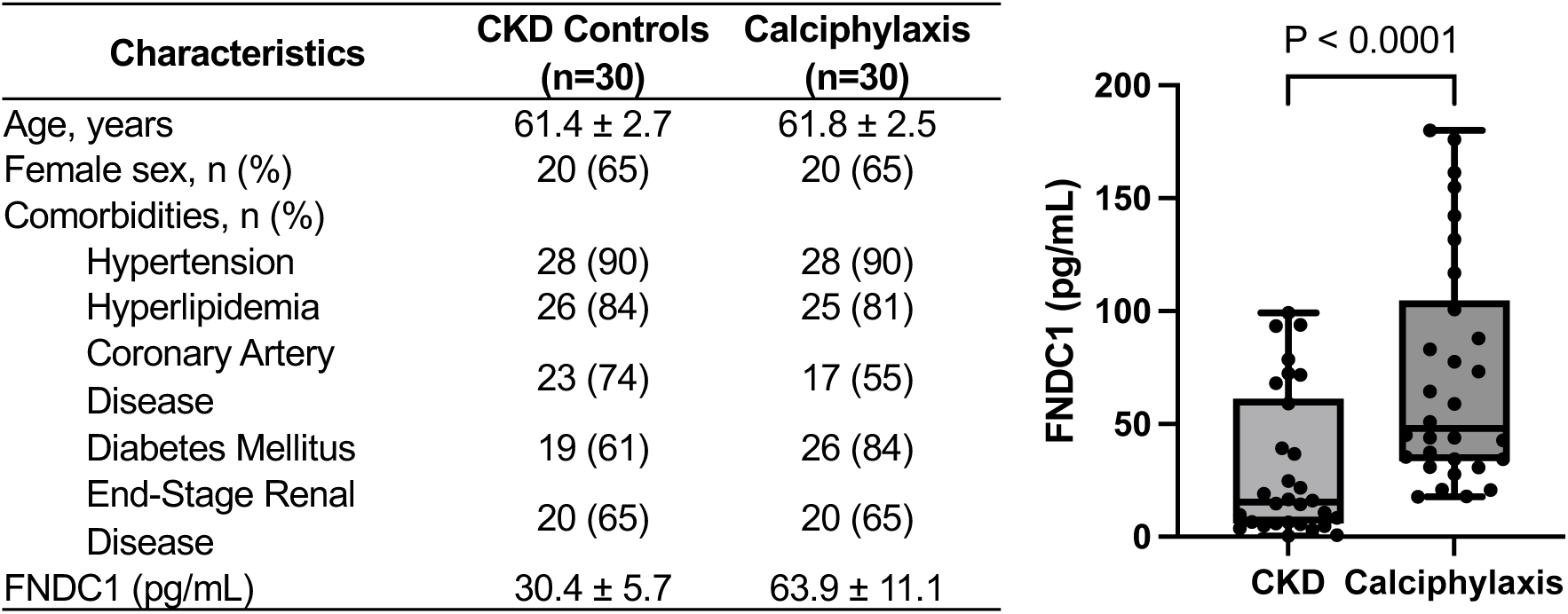
Circulating FNDC1 levels are elevated in calciphylaxis patients compared to matched control patients. **A.** Plasma FNDC1 levels were measured using a sandwich ELISA in 30 calciphylaxis patients and 30 chronic kidney disease (CKD) control patients. Baseline characteristics are provided, revealing similar age, sex, and comorbidities across the two groups. B. On average, plasma FNDC1 was increased more than two-fold in calciphylaxis patients.

To further assess the potential of FNDC1 as a biomarker for coronary artery calcification (CAC), we analyzed the Multi-Ethnic Study of Atherosclerosis (MESA) proteomic cohort, which includes over 3,000 proteomic assays from nearly 2,500 individuals with computed tomography-derived CAC scores. Multivariable linear regression analysis adjusting for age, sex, CKD, diabetes, and hypertension demonstrated that higher FNDC1 levels were independently associated with greater CAC burden (standardized β-coefficient = 0.147 for log[CAC + 1], p = 0.0392; Table 1), demonstrating higher circulating FNDC1 correlated with greater coronary calcification. To determine whether FNDC1 levels predict clinical cardiovascular outcomes, we next examined proteomic and clinical data from 42,687 individuals in the UK Biobank. In covariate-adjusted Cox proportional hazards models, elevated FNDC1 levels were significantly associated with increased risk of myocardial infarction (hazard ratio [HR] 1.21, 95% CI 1.08-1.36, p = 0.0013) and all-cause mortality (HR 1.09, 95% CI 1.02-1.16, p = 0.0064; Table 2), independent of traditional cardiovascular risk factors. Taken together, these results support the utility of FNDC1 as a candidate biomarker for vascular calcification and cardiovascular risk stratification.

**Table 1.** Multivariable Linear Regression Demonstrates Association between FNDC1 and Coronary Artery Calcification After Adjusting for Clinical Risk Factors for Vascular Calcification in the MESA Cohort.

| Model | Parameter | Standardized Coefficient | P-value |
| --- | --- | --- | --- |
| Univariate | FNDC1 | 0.352 | 1.00E-05 |
| Model 1 | Age | 0.113 | 6.23E-94 |
|  | Male Sex | 1.415 | 8.47E-49 |
|  | FNDC1 | 0.199 | 5.17E-03 |
| Model 2 | Age | 0.100 | 4.97E-61 |
|  | Male Sex | 1.376 | 6.35E-47 |
|  | CKD | 0.034 | 6.74E-01 |
|  | DM | 0.403 | 3.74E-05 |
|  | HTN | 0.770 | 3.04E-14 |
|  | FNDC1 | 0.147 | 3.92E-02 |

**Table 2.**
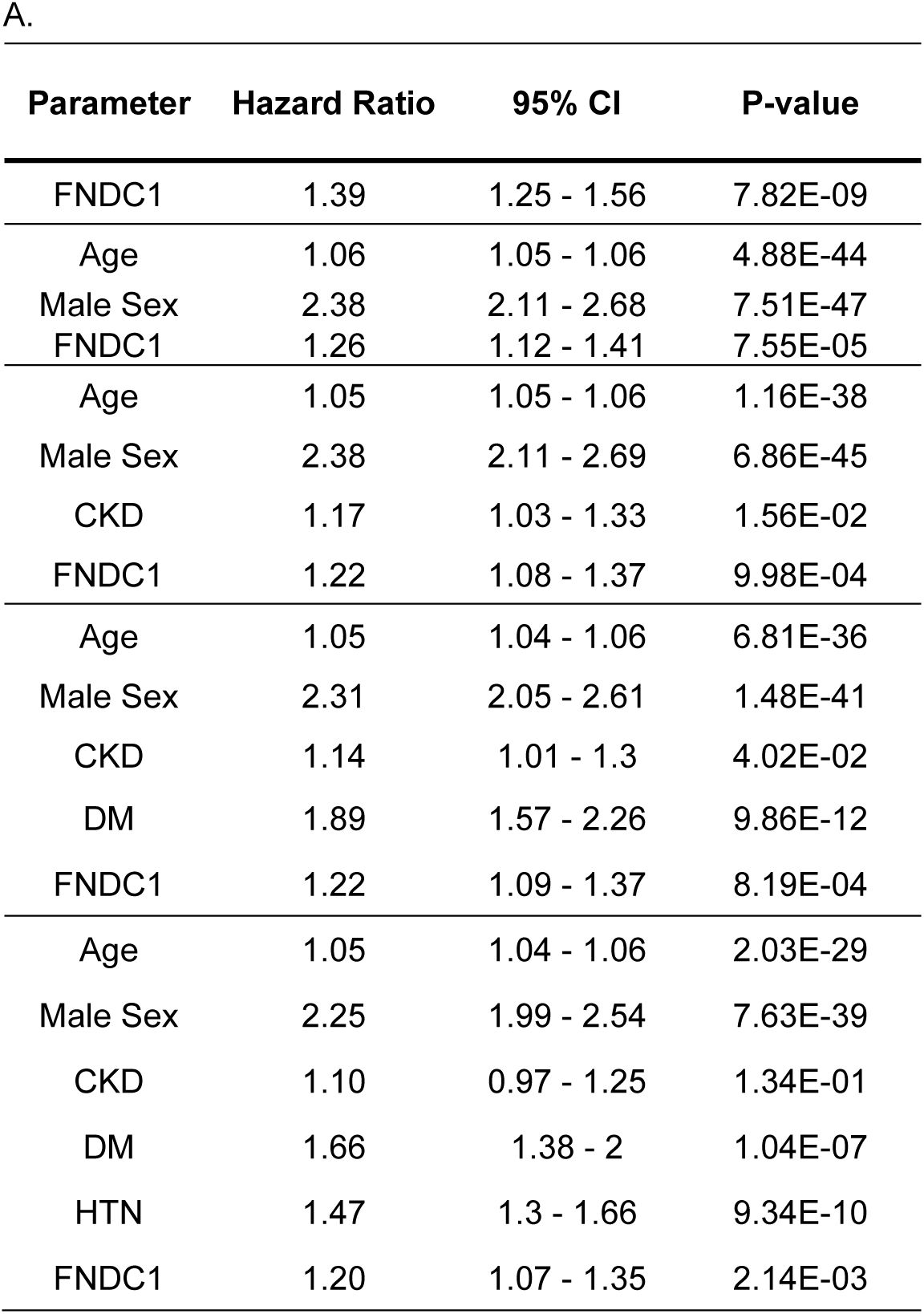
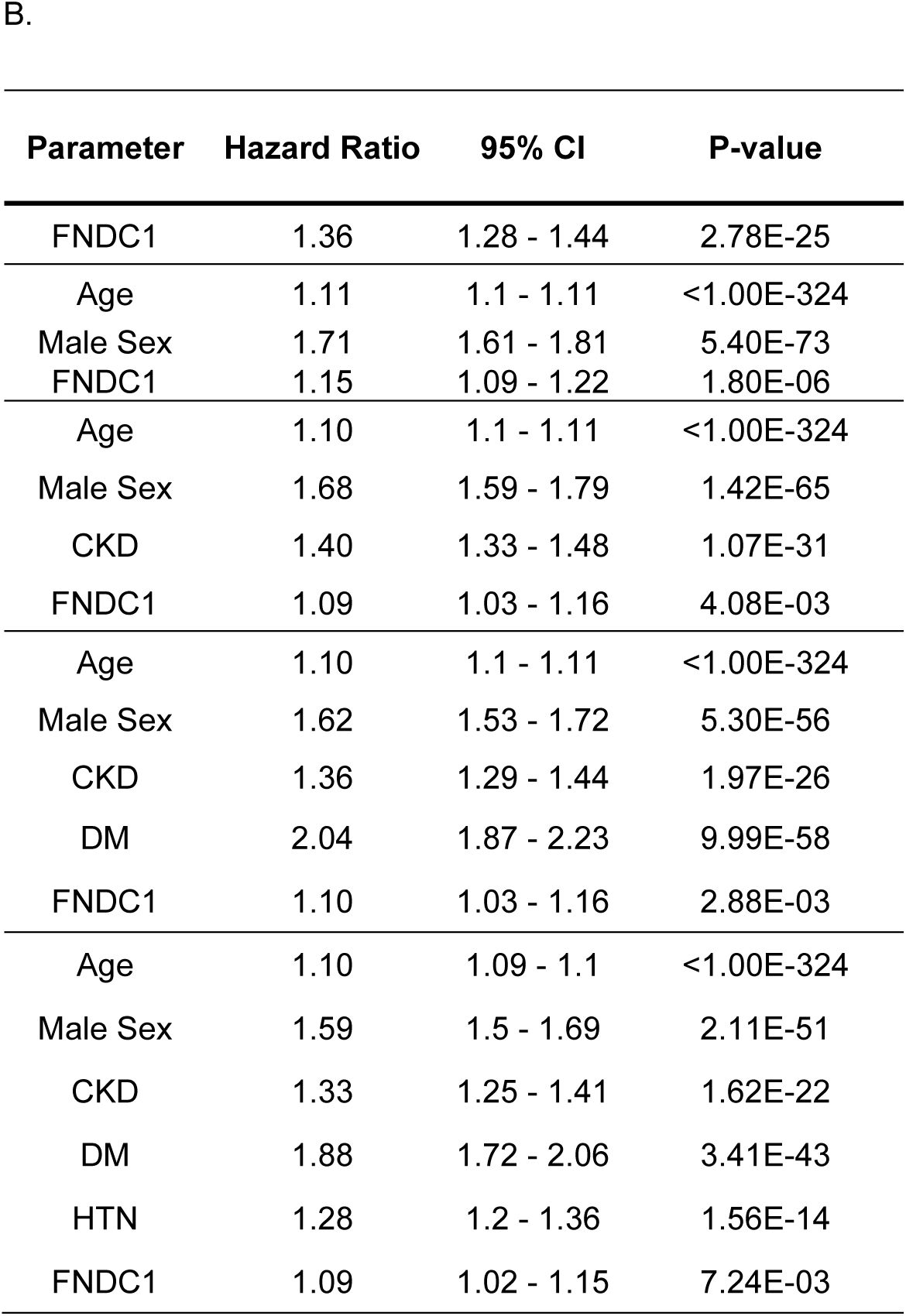
Multivariable Cox Regression Analysis Identifies FNDC1 as a Predictor of (A) Myocardial Infarction and (B) All-Cause Mortality in the UK Biobank (n=42,687)

## DISCUSSION

As the predominant cellular constituent of the arterial wall, VSMCs are essential for maintaining vascular tone and structural integrity under physiological conditions.^50^ However, in response to pathological stimuli, VSMCs exhibit remarkable phenotypic plasticity, acquiring osteogenic-like features that drive medial and intimal calcification in diseases such as atherosclerosis and calciphylaxis.(15, 41) Understanding the molecular mechanisms governing this plasticity remains essential for therapeutic targeting, particularly given the absence of approved treatments for vascular calcification. In this study, we performed the first transcriptomic profiling of human calciphylaxis lesions and identified FNDC1 as a top differentially expressed gene. Although previously implicated in oncogenic signaling, the role of FNDC1 in vascular biology had not been defined. Here, we demonstrate that FNDC1 functions as a potent upstream regulator of VSMC osteogenic reprogramming and vascular calcification. Knockdown and overexpression studies in human vascular smooth muscle cells demonstrated that FNDC1 promotes calcification, proliferation, and migration. Mechanistically, FNDC1-mediated calcification was dependent on activation of the PI3K-AKT signaling axis and upregulation of nicotinamide phosphoribosyltransferase (NAMPT), with pharmacologic inhibition of either pathway attenuating calcification despite FNDC1 overexpression. In two murine models of vascular calcification-Mgp^-/-^ model and vitamin D–induced calcification model-genetic deletion of *Fndc1* conferred protection against aortic calcification. NAMPT inhibition in both models recapitulated the protective effects observed in *Fndc1*-deficient mice. These data position FNDC1 as a novel molecular driver of VSMC calcification and identify AKT and NAMPT as key mediators of FNDC1-induced osteogenic signaling. Beyond its pathogenic role, FNDC1 also exhibited potential as a circulating biomarker. Plasma FNDC1 levels were significantly elevated in patients with calciphylaxis and were independently associated with coronary artery calcification and adverse cardiovascular outcomes in large-scale population cohorts. These findings suggest that FNDC1 may serve as both a mechanistic driver and a clinically relevant biomarker of vascular calcification. Collectively, our results establish FNDC1 as a central regulator of osteogenic reprogramming in VSMCs and nominate the FNDC1–NAMPT-NAD axis as a potential novel therapeutic target in vascular calcification.

FNDC1 belongs to the fibronectin type III domain-containing (FNDC) protein family and contains four highly conserved fibronectin type III (FN3) domains.^51^ FN3 domains are structurally characterized by monomeric β-sandwich folds, which serve as versatile scaffolds for protein-protein interactions and ligand binding.^52–54^ This structural motif underpins the biological activity of multiple FNDC proteins, including FNDC4 and FNDC5, which mediate diverse cellular functions largely through their FN3 domains.^55,56^ Unlike many FNDC family members that are membrane-bound, FNDC1 lacks a hydrophobic transmembrane segment, suggesting its potential to function in both intracellular and secreted forms.^57^ Originally cloned from ischemic rat myocardium, FNDC1 has been shown to regulate hypoxia-induced apoptosis in cardiomyocytes via G protein–coupled signaling pathways.^42,58^ In endothelial cells, FNDC1 has been implicated in vascular endothelial growth factor (VEGF)–mediated angiogenesis, cellular migration, and proliferation.^42^ In cancer models, FNDC1 promotes tumorigenesis through activation of the PI3K-AKT signaling axis, supporting cellular proliferation, invasion, and metastasis.^41,59^ In the current study, we extend the functional repertoire of FNDC1 to the vasculature, demonstrating that FNDC1 promotes osteogenic phenotypic switching of human VSMCs-a process characterized by enhanced proliferation and migration. Mechanistically, this transition is driven by FNDC1-mediated activation of the PI3K-AKT pathway, aligning with its previously characterized role in cancer biology. These findings position FNDC1 as a pleiotropic regulator that integrates growth factor signaling and metabolic reprogramming to promote pathological cellular phenotypes in both tumorigenic and vascular contexts.

While FNDC1 has previously been implicated in the activation of PI3K-AKT signaling, its involvement in cellular metabolic regulation, particularly in the context of NAD⁺ metabolism, has remained uncharacterized. In this study, we uncover a previously unrecognized role for FNDC1 in modulating the NAD⁺ salvage pathway through upregulation of nicotinamide phosphoribosyltransferase (NAMPT), the rate-limiting enzyme responsible for converting nicotinamide (NAM) to nicotinamide mononucleotide (NMN), the immediate precursor for intracellular NAD⁺ biosynthesis. NAD+ is an essential coenzyme that supports fundamental cellular processes including energy metabolism, DNA repair, and cell growth and survival.^46,60^ Given the heightened metabolic demands associated with osteogenic phenotype switch, increased NAD availability may be a requisite for this process.^61,62^ Although the metabolic reprogramming of VSMCs during osteogenic switch remains incompletely characterized, emerging evidence suggests that a shift toward increased glycolytic flux is a key feature of this process.^63^ Because NAD functions as a critical cofactor in glycolysis, our findings raise the possibility that FNDC1-mediated NAMPT induction may facilitate this glycolytic adaptation. The observation that FNDC1 promotes NAMPT expression and NAD^+^ biosynthesis under both normal and osteogenic conditions further underscores its function as a metabolic regulator in VSMCs. Together, these results reveal a novel FNDC1–NAMPT–NAD⁺ axis that may coordinate the metabolic and signaling events necessary for vascular calcification, providing new insight into the molecular underpinnings of VSMC plasticity.

FK866 is a potent and selective small-molecule inhibitor of NAMPT that competitively binds the enzyme’s catalytic domain by structurally mimicking its endogenous substrate, nicotinamide.^61,64^ Through targeted depletion of cytosolic NAD⁺ pools, FK866 has demonstrated robust preclinical efficacy in hematologic and solid malignancies and is currently undergoing evaluation in phase II clinical trials.^47^ Building on these initial findings, next-generation NAMPT inhibitors with enhanced potency and improved pharmacokinetic profiles are actively being pursued to overcome limitations in therapeutic window and tolerability.(55) In light of this growing clinical interest in NAMPT as an oncologic target, the identification of vascular calcification as a previously unrecognized indication for NAMPT inhibition is of particular translational relevance. Our data suggest that repurposing or optimizing NAMPT-directed therapies may represent a viable strategy to modulate vascular osteogenic signaling and mitigate pathological calcification, thereby broadening the therapeutic landscape for cardiovascular disease.

Moreover, the observed associations between elevated circulating FNDC1 levels and the presence of calciphylaxis lesions, increased coronary artery calcification, and adverse cardiovascular outcomes underscore the potential utility of FNDC1 as a novel biomarker of vascular calcification. Currently, calciphylaxis lacks specific diagnostic biomarkers, complicating the early identification of high-risk individuals, even among those with established predisposing risk factors. In this context, incorporation of FNDC1 into clinical assessment frameworks may refine cardiovascular risk prediction and offer mechanistic insights into disease progression. Given its robust association with vascular pathology across diverse cohorts, FNDC1 holds promise not only as a diagnostic biomarker but also as a tool for guiding targeted intervention in patients with multi-vessel calcification.

In summary, we identify FNDC1 as a master regulator of vascular calcification operating through a dual AKT and NAD+ metabolic axis and uncover similarities in molecular dysregulation between small- and large-vessel calcific vasculopathy. Targeting the FNDC1-NAMPT pathway through genetic ablation of FNDC1 and pharmacologic inhibition of NAMPT attenuates vascular calcification *in vivo*, which offers a promising strategy to combat calcific vasculopathies ranging from calciphylaxis to atherosclerosis.

## Acknowledgements

SL is supported by the NIH (F32HL164025). YC is supported by grants from the NIH (HL146103, HL167201 and AG082839), American Heart Association (23TPA1066040), as well as the United States Department of Veterans Affairs (research awards BX005800, CX002706 and BX006321). KR is supported by funding from the National Research Foundation of Korea (RS-2024-00359396). RN is supported by funding from Solaana MD and Pyxis Oncology. RM was supported by the National Heart, Lung, and Blood Institute (R01HL142809 and R01HL162928), the Leducq Foundation, the American Heart Association (22TPA969625), and the Wild Family Foundation.

The MESA projects are conducted and supported by the National Heart, Lung, and Blood Institute (NHLBI) in collaboration with MESA investigators. Support for MESA is provided by contracts 75N92025D00022, 75N92020D00001, HHSN2682015000031, N01-HC-95159, 75N92025D00026, 75N92020D00005, N01-HC-95160, 75N92020D00002, N01-HC-95161, 75N92025D00024, 75N92020D00003, N01-HC-95162, 75N92025D00027, 75N92020D00006, N01-HC-95163, 75N92025D00025, 75N92020D00004, N01-HC-95164, 75N92025D00028, 75N92020D00007, N01-HC-95165, N01-HC-95166, N01-HC-95167, N01-HC-95168, N01-HC-95169, UL1-TR-000040, UL1-TR-001079, UL1-TR-001420, UL1TR001881, DK063491, and R01HL105756. The authors thank the MESA participants and the MESA investigators and staff for their valuable contributions. A full list of participating MESA investigators and institutions can be found at http://www.mesa-nhlbi.org.

## Authors

Sujin Lee (Massachusetts General Hospital)

Yugene Guo (University of Minnesota)

Lova Kajuluri (Massachusetts General Hospital and Harvard Medical School)

Kangsan Roh (Massachusetts General Hospital)

Kuldeep Singh (Massachusetts General Hospital)

Wanlin Jiang (Massachusetts General Hospital, Boston, MA, USA)

Elizabeth Moore (Massachusetts General Hospital)

Helena Tattersfield (Massachusetts General Hospital)

Katrina Ostrom (Massachusetts General Hospital)

Claire Birchenough (Massachusetts General Hospital and Harvard Medical School)

Adam Johnson (Massachusetts General Hospital and Harvard Medical School)

Scott Krinsky (Massachusetts General Hospital)

Houda Bouchouari (Massachusetts General Hospital)

Mohammed Mahamdeh (Massachusetts General Hospital)

Laurel Lee (University of Texas Southwestern Medical Center)

Gregory Wyant (Department of Medicine, Massachusetts General Hospital, Harvard Medical School)

Stephen S. Rich (University of Virginia)

Jerome I. Rotter (The Lundquist Institute for Biomedical Innovation at Harbor-UCLA Medical Center)

Kent Taylor (The Lundquist Institute)

Shaista Malik (University of California, Irvine)

Robert Gerszten (Beth Israel Deaconess Medical Center)

Daniel Cruz (Beth Israel Deaconess Medical Center)

Russell Tracy (University of Vermont)

Hui Wu (Oregon Health and Science University)

Hua Zhang (Integrative Biosciences, School of Dentistry, Oregon Health and Science University)

Haojie Lu (Erasmus MC)

Maryam Kavousi (Erasmus University Medical Center Rotterdam)

Catherine Shanahan (King’s College London)

Melinda Duer (University of Cambridge)

Rafael Kramann (RWTH Aachen University)

Rosalynn Nazarian (Massachusetts General Hospital)

Matthew Eagleton (Massachusetts General Hospital)

Yabing Chen (Oregon Health & Science University)

Clint Miller (University of Virginia)

Sagar Nigwekar (Harvard Medical School)

Rajeev Malhotra (Massachusetts General Hospital and Harvard Medical School)

